# Development and quality assessment of low-cost benchtop malting protocol for laboratory-scale malt quality evaluation

**DOI:** 10.1101/2025.01.01.631005

**Authors:** Heena Rani, Andrew Standish, Jason G Walling, Sarah J Whitcomb

## Abstract

High-quality malt is influenced by three primary factors: barley genotype, environmental conditions, and malting process. To effectively evaluate malting barley breeding material and assess how environmental changes influence malt quality, it is essential to have laboratory- scale malting methods that can produce malt approximating that produced by commercial malting operations. However, existing laboratory-scale malting procedures often demand large quantities of grain, rely on specialized equipment, and are costly. To overcome these challenges, we developed a small sample-scale benchtop malting method utilizing standard laboratory equipment and components available at hardware stores. We validated the method by conducting standard malt quality tests including diastatic power, α-amylase activity, total malt protein, and wort composition (soluble protein, wort soluble/total malt protein, β-glucan, free amino nitrogen, and malt extract). Our findings indicate that the benchtop malting method yields quality metrics comparable to those obtained from established small-scale and full-scale malting protocols. Furthermore, a key innovation of this system is the use of separate Erlenmeyer flasks for malting each sample. Unlike conventional shared malting systems, this design enables precise measurement and comparison of treatment effects across samples malted simultaneously. This reliable, low-cost, and efficient method provides a valuable tool for screening malt quality traits in breeding lines with limited sample sizes and for testing malting regimes aimed at improving malt quality and efficiency. Additionally, it offers an accessible solution for producing high-quality, research-scale malt in laboratories without dedicated quality assurance facilities.

## 1. Introduction

Beer ranks as the third most consumed beverage worldwide, following water and tea. In 2023, global beer production reached approximately 1.88 billion hectoliters, up from 1.3 billion hectoliters in 1998, with China, United States, and Brazil as leading producers (Statista 2024, Accessed on Dec 7, 2024). Beer production is a complex process involving several steps and select raw ingredients including yeasts, hops, and malted barley (Fox & Bettenhausen, 2023). Malt contributes the essential fermentable sugars necessary for fermentation and many of the sensory attributes perceived in beer. Besides its central role in beer production, malt is an ‘intermediate’ in several manufacturing processes such as the production of malt flours, diastatic or non-diastatic malt extracts, cereal syrups, various distilled spirits (such as whiskey, brandy and vodka), and malt vinegar (Bathgate, 2016; Briggs, 1978). The type of malt produced is determined by its intended application.

Malting harnesses the natural germination process to modify hard barley kernels into friable, starch-accessible grains suitable for beer production. This transformation involves the controlled degradation of endosperm cell wall components to expose starch granules and to develop the hydrolytic enzymes whose activity during mashing, a brewing step where malt is crushed and mixed with hot water to produce a sugar-rich fermentable liquid, generates fermentable size sugars from starch chains. The wort provides essential nutrients required for a successful fermentation process. Malting the raw barley comprises three main steps: steeping, germinating and kilning. The objective of steeping is for the grain to rehydrate from an initial ∼12% moisture to 40-45% optimal moisture, soften, and resume metabolism (Almaguer et al., 2024). During the steeping phase, most barley varieties require periodic air rests between water immersion periods to allow the grains to replenish O_2_ and remove CO_2_ (Bettenhausen et al., 2018; Zhang et al., 2017), resulting in a steeping phase of 24-48 h. Once the required moisture content is attained, the grain is then moved into the second phase of malting, germination, where the grain is maintained at a constant temperature and humidity for 4 to 5 days. During germination the grain is turned periodically to 1) increase oxygen and remove CO_2_, 2) to allow the release of heat generated from the germinating grain, and 3) to prevent the rootlets from “matting”. During the germination phase, the hydrolytic and proteolytic enzymes begin degrading cell wall components, a process known as grain modification, which exposes the starch granules in the endosperm for future degradation by amylases during mashing (Carvalho et al., 2016). The fully germinated grain, now referred to as “green malt”, is then dried gradually by a flow of warm air of increasing temperature, called kilning. The goals of kilning are to reduce the moisture content of the green malt to approximately 4-5%, prevent further growth and modification of the grains, to stabilize and preserve the suite of enzymes, and ensure suitable conditions for storage and transportation (Daneri-Castro et al., 2016). Kilning has the most significant influence on malt color and flavor, which depend on factors such as the time, moisture and temperature (Yahya et al., 2014).

Producing high-quality finished malt with optimal enzyme levels, metabolites, and other characteristics is crucial for brewers to maintain constancy and meet the quality goals of their malt beverages. Malt quality is influenced both by characteristics of the procured barley, which are affected by the growing conditions and by physio/biochemical components including husk thickness, starch and protein content, hydrolytic enzyme activity and non-starch polysaccharides, and by the distinct malting conditions applied by the maltster (Gebeyaw, 2021; Yousif & Evans, 2020).

Malt quality analysis entails generation of malt and subjecting it to a battery of lab-based assays from which the quality of the malt can be accurately measured (Turner et al., 2019). The most common quality parameters include total malt extract (a measure of the total fermentable sugars), soluble protein and β-glucan, enzyme activity (diastatic power and α-amylase activity), and wort analyses (viscosity, free amino nitrogen, fine/coarse extract difference, pH and turbidity) (Schmitt & Budde, 2010). Given the complexity of these steps and the need for specialized instrumentation and expertise, the entire process is typically carried out in dedicated malting quality assurance (QA) laboratories.

Breeders of barley for malt production consider malt quality metrics to be critical for varietal development toward commercial release. However, given the cost of QA testing, the limited number of malt QA labs, and slow returns on submitted samples, malt quality testing can be a roadblock in variety development and improvement. Fortunately, simpler, reduced-scale malt quality analysis methods, including isothermal hot water extract of malt (Schmitt et al., 2006) and methods using common laboratory instrumentation for malt and wort analysis (Schmitt & Budde, 2010), have made malt quality assessment more accessible to laboratories lacking dedicated mashing and wort analysis instruments.

Despite these advancements in malt assessment methods, a simplified system to malt small quantities of grain without extensive specialized equipment is still needed. Micro-malting methods have been employed in laboratories for years to support malt research, but these methods typically use 150-500g of raw barley and require specialized equipment and software that are expensive, often requiring an entire room dedicated to housing the equipment. The “tea ball” malting approach (Schmitt & Budde, 2011) offers a small-scale alternative, requiring only 2g of barley, but it still requires specialized micro-malting equipment that can exceed $30k USD making it cost prohibitive for most small scall applications. Commercial malting methods are proprietary and meant for large batch production and therefore not accessible for barley malt varietal screening or malt method testing. While standard malting equipment is available in malt QA labs and other research laboratories devoted to malting research, they are generally not found in most biochemical laboratories due to their substantial size and cost. A more compact benchtop malting apparatus fabricated using general-use laboratory equipment could be attractive to basic researchers interested in studying different aspects of malting.

Here we report the development of a cost-effective benchtop malting apparatus, hereafter “benchtop malting”, that was constructed using standard laboratory equipment. The benchtop malting apparatus and accompanying methods allows for precise control over malting without specialized equipment and is resource-efficient, requiring less water, energy, and space than any design currently available. The benchtop method is easy to implement and requires minimal training, allowing for accessibility of advanced malting research to a broader range of laboratories, as well as educational institutions providing hands-on experience with minimal resource investment. Finally, the benchtop malting method also facilitates rapid prototyping, enabling researchers to quickly test, iterate, and refine malting methods by adjusting key variables such as temperature, time, and moisture levels. This controlled flexibility accelerates innovation and optimization in malting science, making it easier to explore new methods and improvements.

A key advantage of the benchtop malting system over any system developed to date is that each sample is malted in its own closed environment, isolated from the other samples being malted concurrently. In this prototype, the individual barley samples are in separate Erlenmeyer flasks with distinct steep water for each sample, in contrast to the common practice in micro-malting/commercial systems where the same steep water is shared across multiple varieties. This separation allows for greater precision in studying steep-out water composition during different phases of steeping. Furthermore, isolating samples from each other allows for the modulation of environmental conditions and additives, such as minerals, enzymes, growth regulators, and inoculants that can be introduced to individual samples to study their effects on malting without impacting other samples.

We assessed the efficacy of our benchtop malting method by analyzing malt produced from a recommended barley line through standard quality tests. Due to the small sample size and new system, it was necessary to check the reproducibility of the method and compare the quality results with those obtained from standardized malting procedures. Therefore, using the same barley variety, we compared the malt quality results obtained from our benchtop system with those obtained using three other malting methods: 1) micro-malting and 2) tea ball malting, both developed at the Cereal Crops Research Unit as smaller scale, experimental proxies for commercial malting (Fig. S1) as well as 3) commercially produced malt (Rahr Malting Co). The correspondence between the malt quality results obtained from benchtop malting and those from standardized micro-malting and commercial malting supports the conclusion that the combination of benchtop malting and already published reduced-quantity mashing (Schmitt et al., 2006) and malt analysis (Schmitt & Budde, 2010) procedures effectively expand the capacity for preliminary screening of malt quality characteristics.

## 2. Materials and methods

### 2.1. Study design and sample collection

AAC Synergy (Agriculture and Argi-foods Canada), a 2-row, spring-planted malting barley harvested in 2022, was kindly provided to our project by Rahr Malting Co. Before initiating malting, the raw barley seeds were size-sorted and cleaned. Only the seeds retained on a 5.5/64” width slotted screen were malted. The barley was then divided among the four malting experiments, as outlined in section 2.2, with the malting process conducted as described. Each malting method was replicated three times under identical conditions to assess variation among repetitions of the method. The resulting malt was collected after kilning and was gently cleaned by rubbing it against the base of a 1-mm mesh sieve, retaining the kilned malt, while removing the rootlets and other small detritus. Malt quality analysis was performed on nine samples per malting method, three replicates from each of three malting batches.

### 2.2 Malting Processes at different scales

#### 2.2.1 Micro-malting

Micro-malting is the standard malting method employed at the USDA Cereal Crops Research Unit Malt Quality Lab in Madison WI and is used in their barley quality evaluation program. The micro-malting process utilizes three distinct custom-fabricated malting machines designed by Standard Industries (Fargo, ND) to emulate commercial malting conditions, albeit on a smaller scale. These three machines (Fig. S1A-C) operate in series to accomplish the steeping, germination, and kilning stages of malt production (Anderson, 1937; Anderson and Meredith, 1940). The current system capacity allows for generating malts from up to 96 samples of barley at a time, each at a sample size of 100-200 g. Briefly, the steeping process is initiated by immersing cuboidal stainless-steel steeping containers (6” × 3.5” × 3.5”), each with 180 g of sample barley, into the steep water. The steep water is continuously cycled to avoid stagnation and chilled to maintain a temperature of 16°C. The steep phase is 28 h, with 4 h cycles of immersion at 16°C and air rest at 18°C, targeting 45% moisture. The steep-out grains are then transferred to stainless steel cylindrical containers (5.75” diameter × 4” height) with perforated lids for germination at 17°C and >98% humidity for 120 h. The grains are turned every 30 min, with grain moisture adjusted to 45% once during the process. Once germinated, the grains are kilned in cylindrical steel containers (4.25” diameter × 6.5” height) with mesh bottoms following a temperature program of 49°C for 10 h, 54°C for 4 h, 60°C for 3 h, 68°C for 2 h, and 85°C for 3 h with 30 min temperature ramp between each stage. The target final moisture content of the finished malt after kilning is 4%. Micro-malting was conducted in three separate batches, and from each batch three replicate samples from a single container holding 180 g of barley were collected, resulting in a total of nine replicates.

#### 2.2.2 Tea ball malting

“Tea ball” malting was originally developed by Schmitt & Budde (2011) as a means to generate malt from small amounts of grain (< 100 g) (Fig. S1D). Requiring only 3 g of raw grain, each sample is placed in a 3.5 cm diameter stainless steel mesh tea infuser ball (Lund Distribution) and malted in a “Joe White” (previously Joe White Systems, Melbourne Australia) micromalter that can perform each of the three steps (steep, germination and kilning), within a single unit. The tea balls are placed into stainless steel Joe White malting containers, each of which can accommodate 16 tea balls at a time. Carrier barley is added to the container to fill the air pockets between the tea balls to mimic the density of barley grains in a typical germinator. To prepare the malting cans, a base layer of 484 g of carrier barley was added, followed by sixteen tea balls containing the 2 g raw barley grains, and topped with an additional 484 g of carrier barley, creating a sample-to-carrier ratio of approximately 1:28. The contents of the 1 kg containers were steeped to achieve 47% moisture using a 32 h program at 19°C consisting of 8 h wet, 8 h air, 5 h wet, 5 h air, 2 h wet, 2 h air, and a final 2 h wet. Following steeping, the samples were germinated for 48 h at 18°C, then 24 h at 17°C, and an additional 24 h at 16°C. During germination, matted grain was manually separated twice daily at 8 h intervals. At 24 h after initiating the germination program, the boxes containing carrier and sample grains were submerged in water for 45 s, shaken to remove air pockets, and any floating grain were manually submerged. The boxes were then withdrawn, vigorously shaken five times to remove excess water, and then returned to the micromalter to complete germination. Moisture checks were performed at out of steep and after 24, 48, and 96 h of germination. After germination, the tea balls were placed at the bottom of the boxes, covered with carrier grain, and returned to the micromalter for kilning using the same 24 h protocol as mentioned in section 2.2.1. Tea ball malting was also performed in three separate batches. From each batch, material was pooled from 16 tea balls malted in the same stainless steel Joe White malting container. The pooled material was then divided into three replicates, resulting nine total samples (three replicates per malting batch).

#### 2.2.3 Commercial malting

The commercial malt used in this study was kindly provided by Rahr Malting Co (Shakopee MN) and produced using their malting methods designed to generate a standard pale base malt. Due to confidentiality agreements, the details of their protocol cannot be disclosed. Commercial malting was carried out in three batches: the first two batches in Malthouse 2 (using different germination and kiln compartments) and the third batch in Malthouse 5. From each batch, three 30 g samples were collected from the belt after kilning near the beginning, middle, and end of unloading. This material was pooled into a single 90 g composite sample and three replicate samples taken from this composite sample. In total, there were nine commercial malt samples (three replicates per malting batch).

#### 2.2.4 Benchtop malting

The benchtop malting system utilizes inexpensive and commonplace lab equipment as well as parts that are readily available at most retail hardware stores (see parts list below, 2.2.4.1). Several critical factors were considered in developing this experimental malting system to closely approximate conventional malting methods: agitation for germinating grains, airflow and oxygen supply, humidification control.

##### 2.2.4.1 Instrumentation and system components for the benchtop malting system

- shaking water bath (27 l capacity): model SWBR27 (Shel Lab)
- 250 ml Erlenmeyer flask holders (17 units): part #9530531 (Shel Lab)
- refrigerated circulator: model A-25 (Anova), used to maintain precise temperature control of the water bath via recirculating cool water
- custom-fabricated copper manifolds constructed from 3/8” OD copper tubing designed to fit inside the shaking water bath. Used for uniform cooling and conditioned fresh air delivery
- glass aeration wand and 1 l glass storage flask for construction of air humidifier
- aquarium 16-way inline manifold: model a16042900ux0285 (Uxcell), used to distribute humid air to individual barley samples in the system
- 250 ml Erlenmeyer flasks for water trap (1 unit) and to contain barley samples for steeping (16 units) and germination (16 units)
- #6 drilled 2-hole stoppers (2 units) for construction of air humidifier and water trap
- vinyl tubing (3/8” ID x 1/2” OD) to connect components of the malting system such as the copper manifolds and air humidifier
- silicone aquarium tubing (4 mm) to connect the inline manifold to the individual 250 ml Erlenmeyer flasks containing the germinating barley samples
- zip ties and hose clamps to hold the components of the malting system in place
- programmable digital timer: Model T319 (Nearpow), used to control the periodic shaking of the water bath
- programmable oven: Thermocenter TC 40S (Salvis), used to control the kilning temperature regime including 30 min ramps between different kilning stage temperatures.
- non-programmable oven: Heratherm OMS60 (Thermo Scientific), used to control the kilning temperature regime without the ability to implement 30-minute ramps between different kilning stage temperatures.

##### 2.2.4.2 Instrumentation assembly

Refer to Fig. 1a and 1b for assembly details.

1. The refrigerated circulator inlet and outlet were connected to one of the custom-fabricated copper manifolds with vinyl tubing. The copper manifold was placed along the inside perimeter of the shaker bath shelf and was secured to the shelf with zip ties.
2. The air humidifier was assembled from a 1 l glass storage flask fitted with a pre-drilled #6 2-hole stopper. A glass aeration wand was inserted through one of the holes in the stopper and positioned inside the flask. Using vinyl tubing, the air humidifier inlet was connected to a source of compressed air, and the outlet was connected to the inlet of the other custom-fabricated copper manifold. The copper manifold was placed along the inside perimeter of the shaker bath shelf and was secured to the shelf using zip ties.
3. The outlet of the conditioned air copper manifold was connected to the inlet of a water trap constructed from a 250 ml Erlenmeyer flask fitted with a pre-drilled #6 2-hole stopper. The water trap outlet was connected with vinyl tubing to the inlet of the aquarium 16-way inline manifold. The aquarium manifold was secured to the cross bar of the shaker bath shelf using zip ties.
4. The water trap and 250 ml Erlenmeyer flasks used to contain the developing barley malt were secured to the shelf of the shaker bath using the flask holders for the shaker.
5. The outlet ports of the aquarium manifold were connected to lengths of silicone aquarium tubing. The free ends of the tubing were positioned in the bottoms of the flasks used to contain the developing barley malt.
6. The shaking water bath was connected to the programmable timer during germination.

**Fig. 1.**
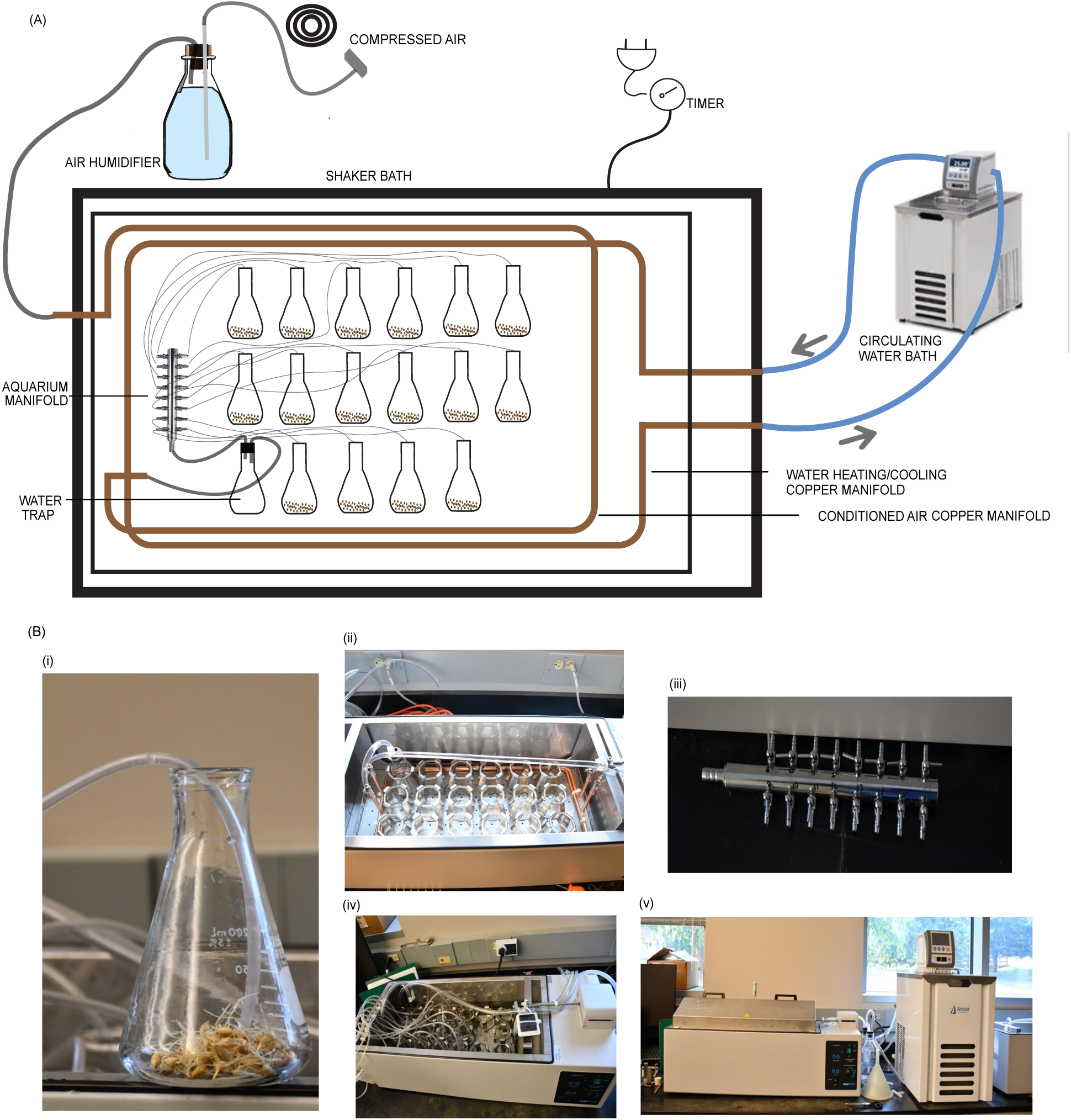
Overview of the benchtop malting system. (A) Schematic diagram; (B) Photos of the apparatus: (i) germinating seeds using the benchtop system, (ii) copper manifold setup within a water bath for cooling water and delivering humid air, (iii) aquarium 16-way inline manifold, (iv) silicone aquarium tubing connecting the manifold to individual 250 mL Erlenmeyer flasks, (v) water bath for malting and the refrigerated circulator for maintaining precise temperature control via recirculating cool water.

##### 2.2.4.3 Benchtop malting protocol

The steeping process was conducted in 250 ml Erlenmeyer flasks, where 3 g of seeds were soaked in 50 ml of cold tap water for 32 h at 19°C, following a schedule of 8 h wet, 18 h air rest, and 6 h wet. The moisture content of the seeds during steeping was maintained at 48- 48.5%. It was observed that maintaining the seed moisture content slightly above 47% during steeping minimizes the need for further moisture adjustment during germination, as seeds typically lose approximately 1% moisture during this phase. In cases where the moisture content of the steep-out seeds exceeded the target level, excess moisture was removed using Kimwipes. The steeped seeds were then transferred to fresh, dry Erlenmeyer flasks for germination. The germination process was carried out for 48 h at 18°C, followed by 24 h at 17°C, and an additional 24 h at 16°C. Dematting and moisture check were performed manually after 60 h (on the third day of germination). Following germination, the seeds were kilned using two distinct protocols: one in a programmable oven and the other in a non-programmable oven. The programmable oven allowed slower temperature ramps between kilning stages, enabling a gradual increase in temperature and adjustable fan speed (we used 60% of its maximum speed) to prevent excessive drying of the seeds. The kilning steps were as follows: from 28°C to 49°C at 0.7°C/min followed by 10 h at 49°C, from 49°C to 54°C at 0.2°C/min followed by 4 h at 54°C, from 54°C to 60°C at 0.3°C/min followed by 3 h at 60°C, from 60°C to 68°C at 0.3°C/min followed by 2 h at 68°C, and from 68°C to 85°C at 0.6°C/min followed by 3 h at 85°C. In the non-programmable oven, the same final temperatures and holding durations were achieved; however, the temperature transitions between steps (49°C to 54°C, 54°C to 60°C, 60°C to 68°C, and 68°C to 85°C) occurred at a fixed ramp rate of 2°C per minute, as the oven lacked the capability for programmable ramps (Fig. S2). Despite this limitation, the non-programmable oven was tested because of its widespread availability in research laboratories. For each benchtop malting batch, the material in three flasks was pooled to create a composite sample. The pooled sample was then divided into three replicates, resulting in nine total samples (three replicates per malting batch).

### 2.5 Malt quality analysis

Malt grinding and mashing were performed following the protocols described in Schmitt et al., 2006 and Schmitt & Budde, 2010. Small-scale malt quality analysis was conducted using the methodology described in Schmitt & Budde, 2011.

### 2.6 Statistical analysis

One-way ANOVA was performed for each malt quality parameter to assess differences among malting processes (micro-malting, commercial, tea ball, benchtop method using programmable oven and benchtop method using non-programmable oven). Post-hoc analysis using Tukey’s Honest Significant Difference (HSD) test was applied to identify significant differences between malting methods. Compact letter display (CLD) was used to group similar treatments for easier interpretation. Analyses were conducted in R, using the multcomp (v1.4.25) (Hothorn et al., 2008) and multcompView (v0.1.10) (Graves et al., 2019) packages for post-hoc analysis and CLD visualization, with a significance level of 0.05.

## 3. Results and discussion

The aim of this study was to develop a benchtop malting apparatus that offers a simple and cost-effective setup for producing high-quality malt from small quantities of grain, comparable to malt generated by larger and more expensive systems commonly employed in research and industrial malt quality laboratories. To closely mimic conventionally accepted experimental malting methods, several essential features were incorporated. Periodic agitation was introduced during germination to mitigate gravitropic effects and prevent rootlet matting. Fresh air circulation was maintained during the germination phase to remove carbon dioxide and supply oxygen essential for the respiration of barley kernels. This airflow also helped to minimize microbial growth during malting. Humidification control was carefully managed throughout the steeping and germination phases to achieve proper kernel modification and avoid the production of under-modified malt. Furthermore, a key innovation of this system is the use of separate Erlenmeyer flasks for malting each sample. Unlike conventional shared malting systems, this design enables manipulation of individual malting samples independently from others being malted concurrently under the same malting regime.

To evaluate the benchtop system’s performance in producing malt of comparable quality to several standardized malting methods (as described in section 2.2), various malt quality parameters were measured and compared. These included malt extract (ME), diastatic power (DP), and α-amylase activity, as well as wort composition parameters such as soluble protein, soluble/total protein ratio (S/T), β-glucan, and free amino nitrogen (FAN). The results from these analyses are presented in Figs. 2 and 3.

**Fig 2.**
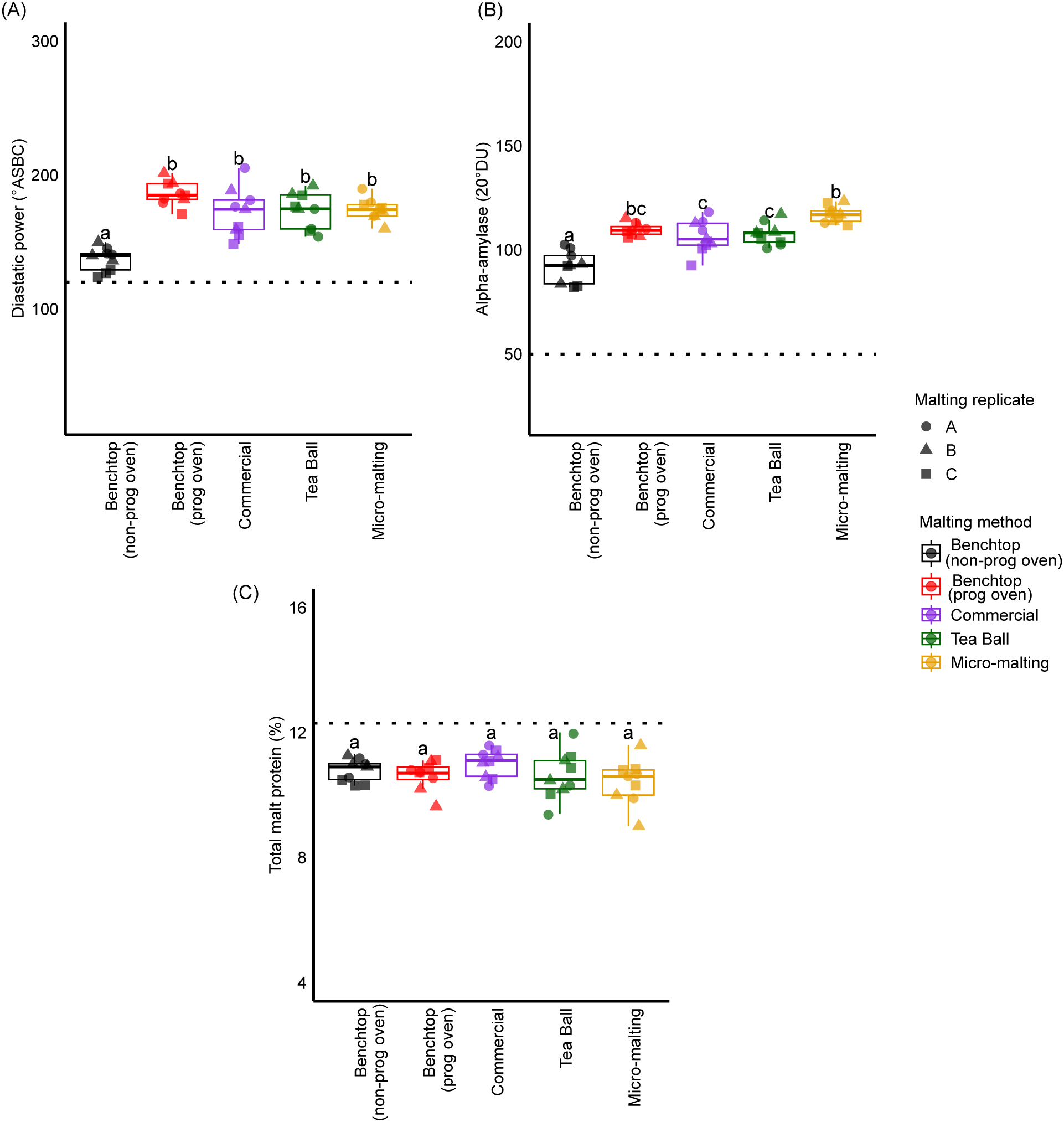
Comparison of malt parameters across different malting methods. For each malting method, measured values from each of the nine samples (n = 3 per malting replicate) are shown as points and the shape of the point corresponds to the malting replicate. The box plots show the median and interquartile range of those values. Letters above the boxes indicate groupings determined by Tukey’s HSD test, where methods sharing the same letter are not significantly different (p <0.05). Horizontal black dashed lines represent criteria for evaluating malt quality based on the American Malting Barley Association (AMBA) guidelines; these include diastatic power (>120 °ASBC), alpha-amylase (>50, 20°DU), and malt protein (≤12.8%).

**Fig 3.**
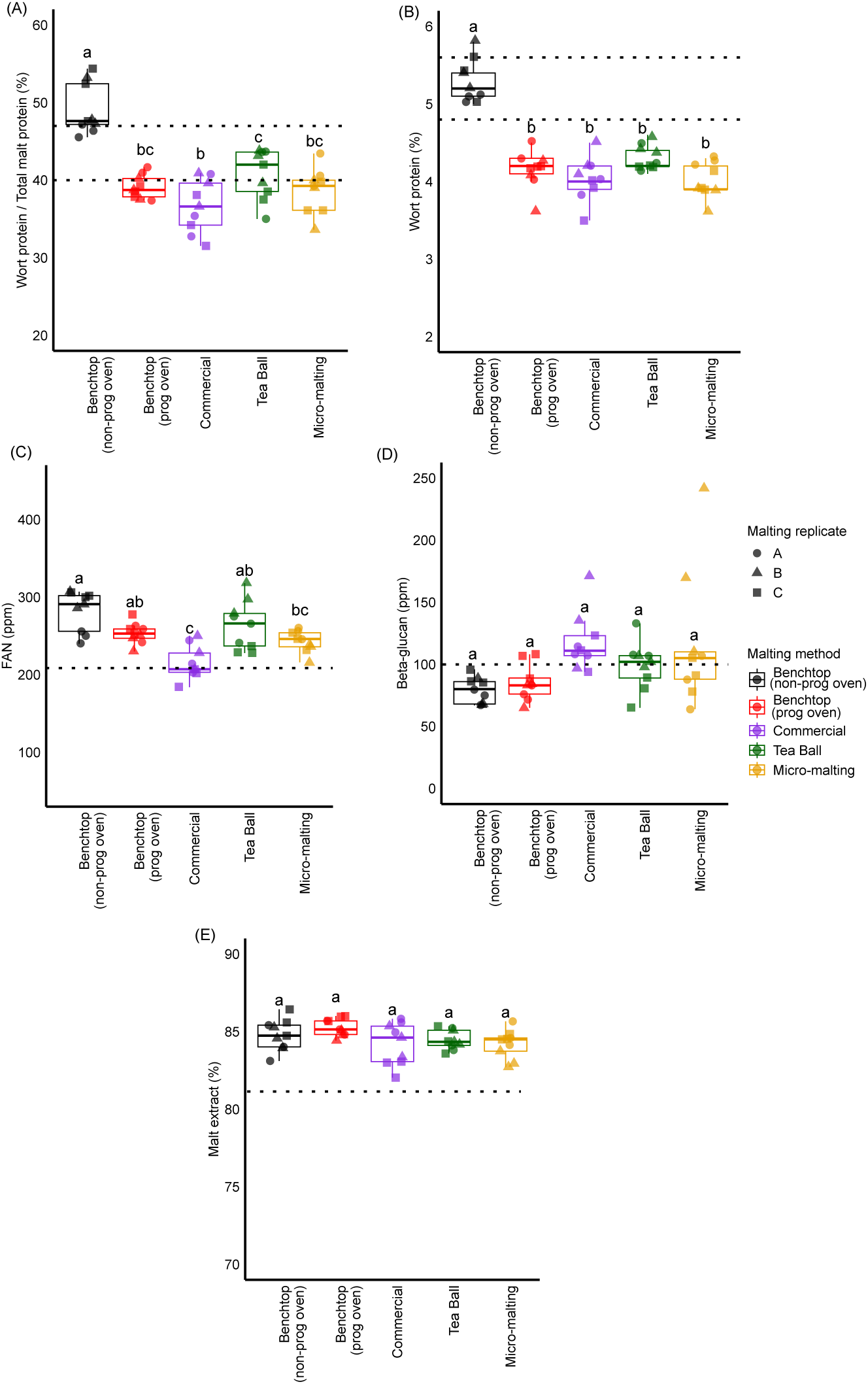
Comparison of wort extract parameters across different malting methods. For each malting method, measured values from each of the nine samples (n = 3 per malting replicate) are shown as points and the shape of the point corresponds to the malting replicate. The box plots show the median and interquartile range of those values. Letters above the boxes indicate groupings determined by Tukey’s HSD test, where methods sharing the same letter are not significantly different (p <0.05). Horizontal black dashed lines represent criteria for evaluating malt quality based on the American Malting Barley Association (AMBA) guidelines; these include wort protein/total protein (40-47%), wort protein (4.8-5.6%), FAN (>210 ppm), beta-glucan (<100 ppm), malt extract (>81%).

### 3.1 Malt quality analysis for benchtop malting system and standard malting methods

#### 3.1.1 Diastatic power (DP) and α-amylase activity

The DP of barley malt reflects the combined activity of key starch degrading enzymes, namely α-amylase, β-amylase, limit dextrinase, and α-glucosidase (Gibson et al., 1995; Walker & Panozzo, 2016). These enzymes collectively facilitate the breakdown of starch into fermentable sugars, with α-amylase initiating the process by hydrolyzing α-1,4-glycosidic linkages, followed by the exo-acting β-amylase cleaving maltose units from the non-reducing ends of starch molecules. Limit dextrinase and α-glucosidase further contribute to the efficient degradation of starch by attacking branched structures and producing glucose, respectively (Evans et al., 2009). Moderate to high DP is a desirable malt characteristic because it supports efficient starch-to-sugar conversion and this generally correlates well with increased ME yields (Henson & Duke, 2007). In the current study, DP and α-amylase activity in the malt produced by the benchtop system and kilned in a programmable oven was not significantly different from that in the malt produced by the other malting methods (Fig. 2A). Initially, the DP and α-amylase activity measurements from the benchtop were significantly lower (1.3-fold and 1.2-fold, respectively) than the other two methods and the use of a non-programmable oven for kilning green malt (Fig. 2A-B) was identified as the likely cause. Malt kilned in the non-programmable oven experienced faster temperature ramps between holding temperatures and high fan speed, both of which may have contributed to accelerated moisture loss and enzyme denaturation, and thereby lower DP and α-amylase activity. Additionally, β-amylase, being more heat-sensitive than α-amylase (De Schepper et al., 2022; Evans et al., 2003), is likely to denature even more rapidly under such conditions, further contributing to the reduction in overall DP.

#### 3.1.2 Protein content and S/T ratio

Total protein levels showed no significant differences among malts produced by different methods, including benchtop malting using the non-programmable oven (Fig. 2C). This suggests that the absence of temperature ramps during kilning did not impact the overall total protein content in the malt, since total malt protein content is primarily determined by intrinsic factors such as the barley grain itself, soil fertility, and environmental factors rather the kilning process (Karababa et al., 1993).

However, the S/T ratio was affected by different kilning methods, with the non-programmable oven producing malt with significantly higher S/T protein than other methods (Fig. 3A). The elevated S/T ratio in this case can be explained by the higher wort soluble protein content from malt produced with the non-programmable oven (Fig. 3B). This increase in soluble protein suggests that the non-programmable oven may have caused more extensive protein degradation during kilning, leading to the formation of smaller protein fragments, which are more soluble in wort, thereby raising the S/T ratio.

#### 3.1.3 Wort free amino nitrogen (FAN)

FAN is defined as the sum of the individual amino acids, ammonium ions, and small peptides (di- and tripeptides) in wort (Hill & Stewart, 2019). A significant portion of FAN is generated during steeping in the malting process, while the remaining nitrogenous compounds are produced by protease activity during mashing of malt (Lei et al., 2013). In this study, no significant differences in FAN levels were observed between malt produced by the benchtop system and other malting systems, except for the commercial malt, which exhibited significantly lower FAN levels (Fig. 3C). These results indicate that benchtop malting systems, regardless of whether programmable or non-programmable ovens are used for kilning, produce malt with similar FAN levels to other more standardized laboratory malting methods. The lower FAN content in commercially malted barley could be attributed to factors such as the use of gibberellic acid (GA) during germination, which was not used in other malting methods tested in this study.

#### 3.1.4 **β**-glucan content

β-glucan is a major polysaccharide found in barley cell walls, and its breakdown during malting is crucial to avoid issues such as high wort viscosity and filtration problems during brewing (Rani et al., 2024b). The initial concentrations of β-glucan in barley and malt can serve as reliable predictors of their levels in wort (Habschied et al., 2020). The β-glucan levels in wort produced by benchtop malting systems, whether using programmable or non-programmable ovens for kilning, were not significantly different from the other malting methods tested (Fig. 3D).

Unlike DP and α-amylase activity, the absence of temperature ramps in the non-programmable oven did not affect wort β-glucan levels.

#### 3.1.5 Malt extract (ME)

ME, which represents the quantity of dissolved solids derived from malt, is a critical factor for maltsters and brewers when evaluating or selecting malting barley. The extract yield directly affects the efficiency of beer production and is therefore arguably the largest economic factor of the all quality parameters measured. High extract levels are also a key objective in breeding programs aimed at developing superior malting cultivars (Li et al., 2008). In this study, no significant differences in ME were observed across the methods tested (Fig. 3E) suggesting that the benchtop malting method produces malt with comparable extract potential to established malting methods. These results highlight the reliability of the benchtop method for assessing ME and its potential application in breeding and research purposes.

### 3.2 Reproducibility among malting replicates in benchtop and standard malting systems

To evaluate the reproducibility of malt quality parameters across different malting systems, the coefficient of variation (CV) was analyzed. CV provides a measure of relative consistency across multiple malting runs, facilitating the comparison of variability in parameter measurements.

Overall, CV values for most malt quality parameters remained consistently low (<12.5%), reflecting reproducible measurements across malt replicates from both the benchtop and standard methods. However, wort β-glucan exhibited the highest variability among the measured attributes. Specifically, the variation in wort β-glucan measurements among replicates was highest in the micro-malting system (42%). These results align with prior observations that wort β-glucan measurements are among the most variable routine quality assurance metrics (Schmitt & Budde, 2011). However, the benchtop malting system demonstrated improved reproducibility for β-glucan, with markedly lower CV values (Fig. 4). These findings underscore the robustness of the benchtop malting system in generating reproducible data for malt quality parameters.

**Fig 4.**
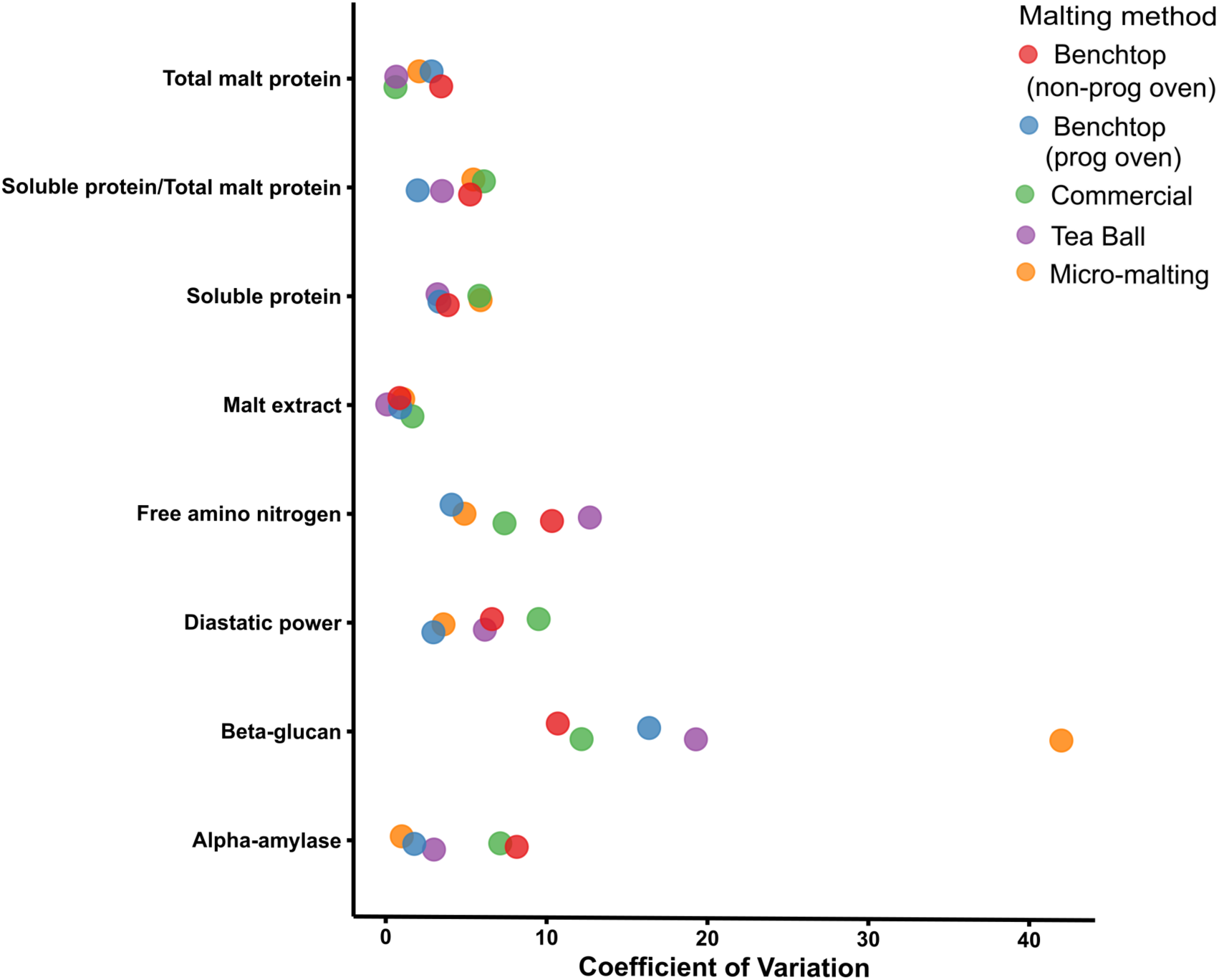
Distribution of coefficient of variation (CV) values for malt quality parameters across different malting methods. The data points represent the CV values for each parameter and are colored by malting method.

## 4. Conclusion

This study demonstrated that our benchtop malting system can produce malt with quality comparable to established methods, such as micro-malting, tea ball malting, and commercial malting. The consistency of key malt quality parameters across these methods validates the benchtop system as a reliable tool for small-scale malting research. Additionally, our findings highlight the importance of controlled temperature ramps during kilning, as the use of a non-programmable oven resulted in a significant reduction in DP and -amylase activity, an increase in the S/T ratio, and underscores the importance of gradual heating during kilning for preserving enzyme activity and enhancing malt quality. However, while the non-programmable oven produced significant differences in some parameters, the magnitude of these differences were not large. Thus, the non-programmable oven can still serve as a practical alternative for kilning, especially for parameters like β-glucan content, FAN, total malt protein, and ME.

In ongoing research, we are further evaluating the suitability of the benchtop malting system for screening breeding material with varying malt characteristics. Based on the results presented above, we expect the benchtop malting system to effectively distinguish between low, medium, and high-quality malting barley material from breeding programs. If so, this will give breeders and researchers a reliable, low-cost, and efficient option for screening malt quality traits with limited sample quantity. Additionally, it offers an affordable solution for testing malting regimes to improve malt quality and efficiency.

## Supporting information

Supplementary Fig S1 ans S2

## Acknowledgements

The authors wish to express their gratitude to Frankie Maynard, Leslie Zalapa, and Chris Martens for their invaluable contributions to this study. Frankie Maynard assisted with sample weighing and grinding for malt quality analysis. Leslie Zalapa provided guidance in Tea Ball malting and malt quality analysis. The micro-malting experiment was performed by Chris Martens at the USDA-ARS Cereal Crops Research Unit, Madison, WI. The authors also extend their thanks to Rahr Malting Co. for providing commercially malted samples, which greatly contributed to the scope and quality of this research.

## 6. CRediT authorship contribution statement

**Heena Rani** - conceptualization, methodology, investigation, data curation, formal analysis, validation, visualization, and writing – original draft. **Andrew Standish** - methodology, investigation, validation, visualization and writing – original draft. **Jason G Walling**- writing – review and editing, supervision, funding acquisition, resources, project administration, and validation. **Sarah J Whitcomb** - conceptualization, validation, writing – review and editing, supervision, funding acquisition, resources, and project administration.

## 7. Funding

This research was supported in part by an appointment to the Agricultural Research Service (ARS) Research Participation Program administered by the Oak Ridge Institute for Science and Education (ORISE) through an interagency agreement between the U.S. Department of Energy (DOE) and the U.S. Department of Agriculture (USDA). ORISE is managed by ORAU under DOE contract number DE-SC0014664. All opinions expressed in this paper are the author’s and do not necessarily reflect the policies and views of USDA, DOE, or ORAU/ORISE.

Mention of trade names or commercial products in this publication is solely for the purpose of providing specific information and does not imply recommendation or endorsement by the U.S. Department of Agriculture. USDA is an equal opportunity provider and employer.

