## Supplementary Fig S1 ans S2 for "Development and quality assessment of low-cost benchtop malting protocol for laboratory-scale malt quality evaluation"

(A)

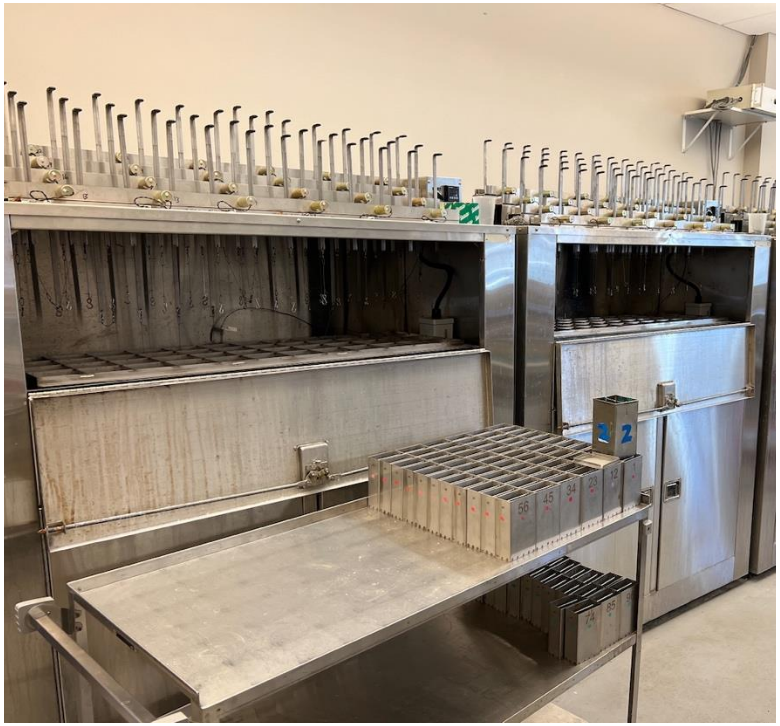

(B)

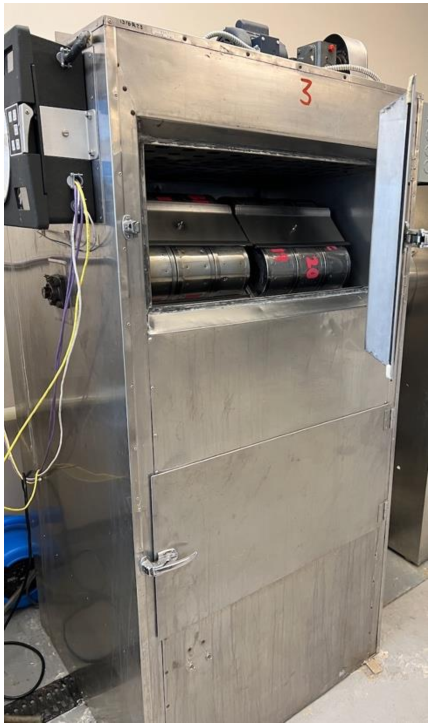

(C)

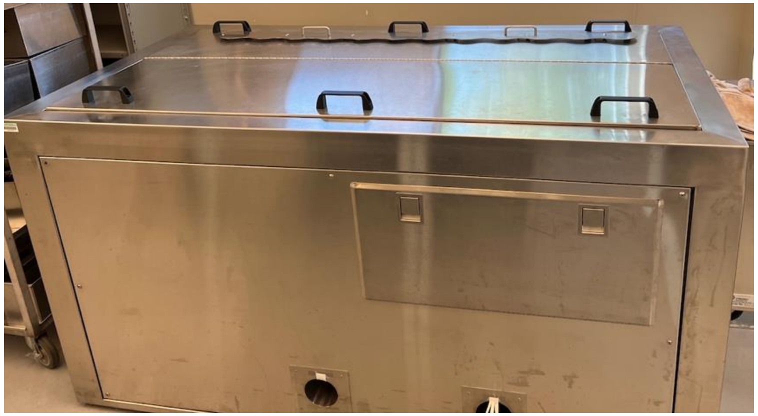

(D)

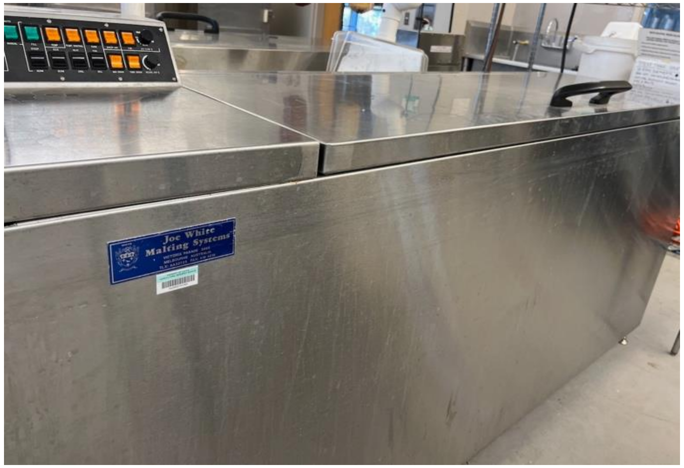

Fig. S1: Micro-malting and Joe White malting systems at the Cereal Crops Research Unit, serving as smaller-scale experimental proxies for commercial malting processes. (A) Steep tanks used in the micro-malting system, (B) Germination tank for the micro-malting system, (C) Kilning machine for the micro-malting system, and (D) Joe White malting system utilized for tea ball malting.

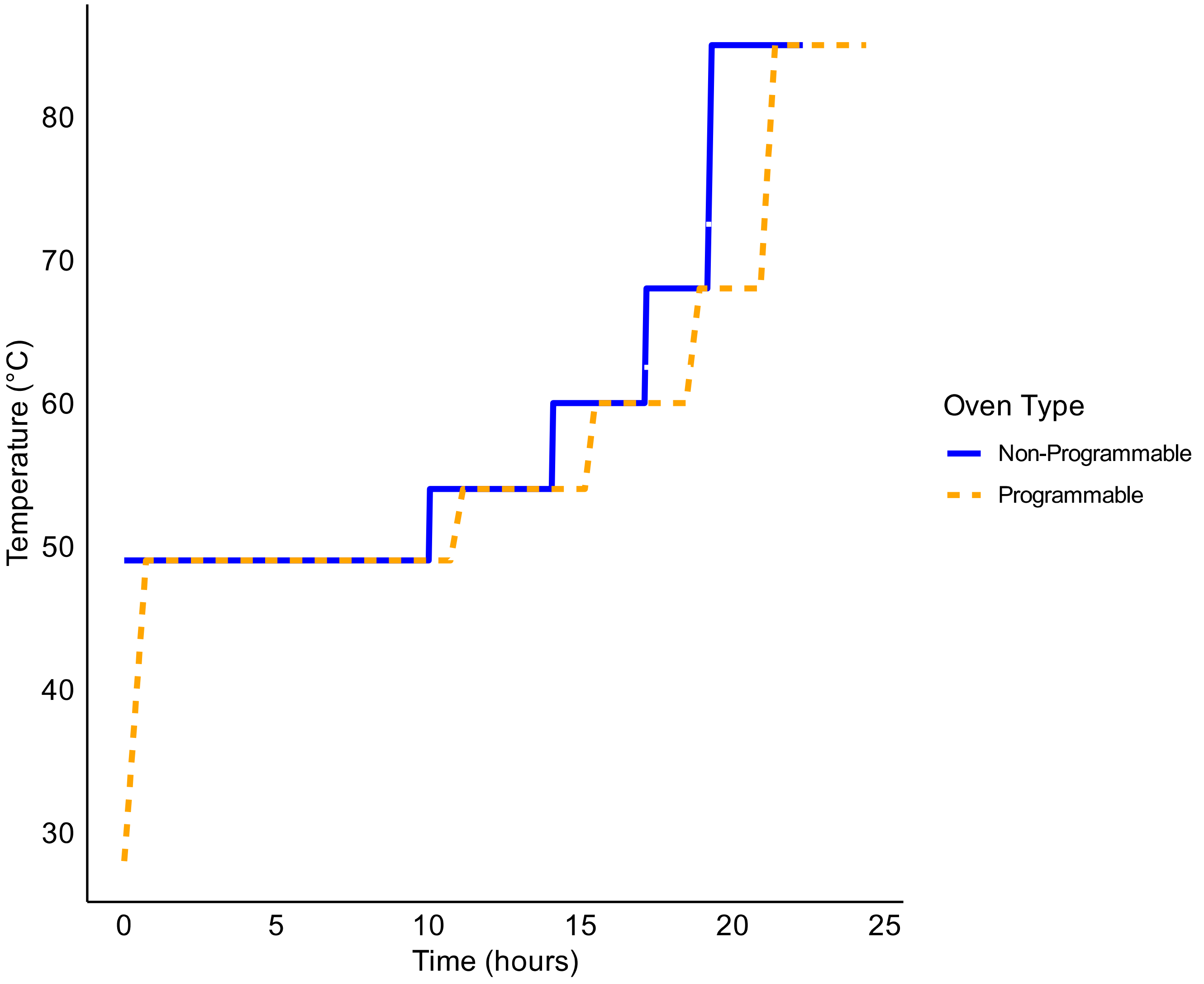


Fig. S2: Comparison of kilning temperature profiles between programmable and non-programmable ovens
